# metaRange: A framework to build mechanistic range models

**DOI:** 10.1101/2024.03.07.583922

**Authors:** Stefan Fallert, Lea Li, Juliano Sarmento Cabral

## Abstract

1. Mechanistic or process-based models offer great insights into the range dynamics of species facing non-equilibrium conditions, such as climate and land-use changes or invasive species. Their consideration of underlying mechanisms relaxes the species-environment equilibrium assumed by correlative approaches, while also generating conservation-relevant indicators, such as range-wide abundance time-series and migration rates if demographically explicit. However, the computational complexity of mechanistic models limits their development and applicability to large spatiotemporal extents.
2. We developed the R package ‘metaRange’ that is a modular framework to build population-based and metabolically constrained range models. It provides a function catalogue for users to calculate niche-based suitability, metabolic scaling, population dynamics, biotic interactions, and kernel-based dispersal, which may include directed movement. The framework’s modularity enables the user to combine, extend, or replace these functions, making it possible to customize the model to the ecology of the study system. The package supports an unlimited number of static or dynamic environmental factors as input, including climate and land use.
3. As an example, we simulated 100 virtual species in Germany on a 1 km^2^ resolution over 110 years under realistic environmental fluctuations in three scenarios: without competition, with competition, and with competition and a generalist-specialist trade-off. The results are in accordance with theoretical expectations. Due to the population-level, the package can execute such extensive simulation experiments on regular end-user hardware in a short amount of time. We provide detailed technical documentation, both for the individual functions in the package as well as instructions on how to set up different types of model structures and experimental designs.
4. The metaRange framework enables process-based simulations of range dynamic of multiple interacting species on a high resolution and low computational demand. We believe that it allows for theoretical insights and hypotheses testing about future range dynamics of real-world species, which may better support conservation policies targeting biodiversity loss mitigation.

## 2 Introduction

The ongoing human-induced environmental change is one of the biggest challenges for humanity and for all the other species living on this planet (IPBES, 2019). Models are needed that can reliably predict how impending environmental change affects the dynamics of occurrence, populations, and range of species (Cabral et al., 2017; IPBES, 2019; Urban et al., 2022, 2016).

The widely available and used phenomenological species distribution models (SDMs) can achieve a high accuracy when predicting present day habitat suitability, but because they assume species-environment equilibrium and there is no direct mechanistic links between the predictor variables and the estimated suitability, spatiotemporal extrapolations are problematic (Briscoe et al., 2019; Dormann et al., 2012). Mechanistic or process-based models relax equilibrium assumptions by explicitly accounting underlying causality while, besides habitat suitability, also predicting individual survival, population sizes, and/or species richness depending on integrated mechanisms (Cabral et al., 2017). However, these models are difficult to develop and apply because of the theoretical underpinnings, computational complexity and demand, and the sparse data availability for calibration (Briscoe et al., 2019; Dormann et al., 2012; Zurell et al., 2016). Consequently, most human-impact assessments still use the classic phenomenological SDMs (Zurell et al., 2022; but see Cabral et al., 2013, 2011), despite urgent calls for mechanistic assessments (IPBES, 2019).

To answer those calls, we developed a modular framework to build process-based range models. Because our framework is population-based, it can use a small number of free parameters compared to individual-based models. Due to the modularity, users can build mechanistic models that can include various processes such as reproduction, dispersal, species interactions and metabolic constraints. This framework is implemented as an R package, allowing for simulations on a high resolution, with large range-wide extents, and high number of interacting species.

## 3 Package Description

The R package metaRange is a tool to build mechanistic range models that are focused on population and metapopulation dynamics of one or multiple interacting species. A key feature of metaRange is the extensibility that emerges from its object-oriented implementation.

The package is built around the concept of a ‘simulation’ object that manages the simulation state (Fig. 1).

**Figure 1:**
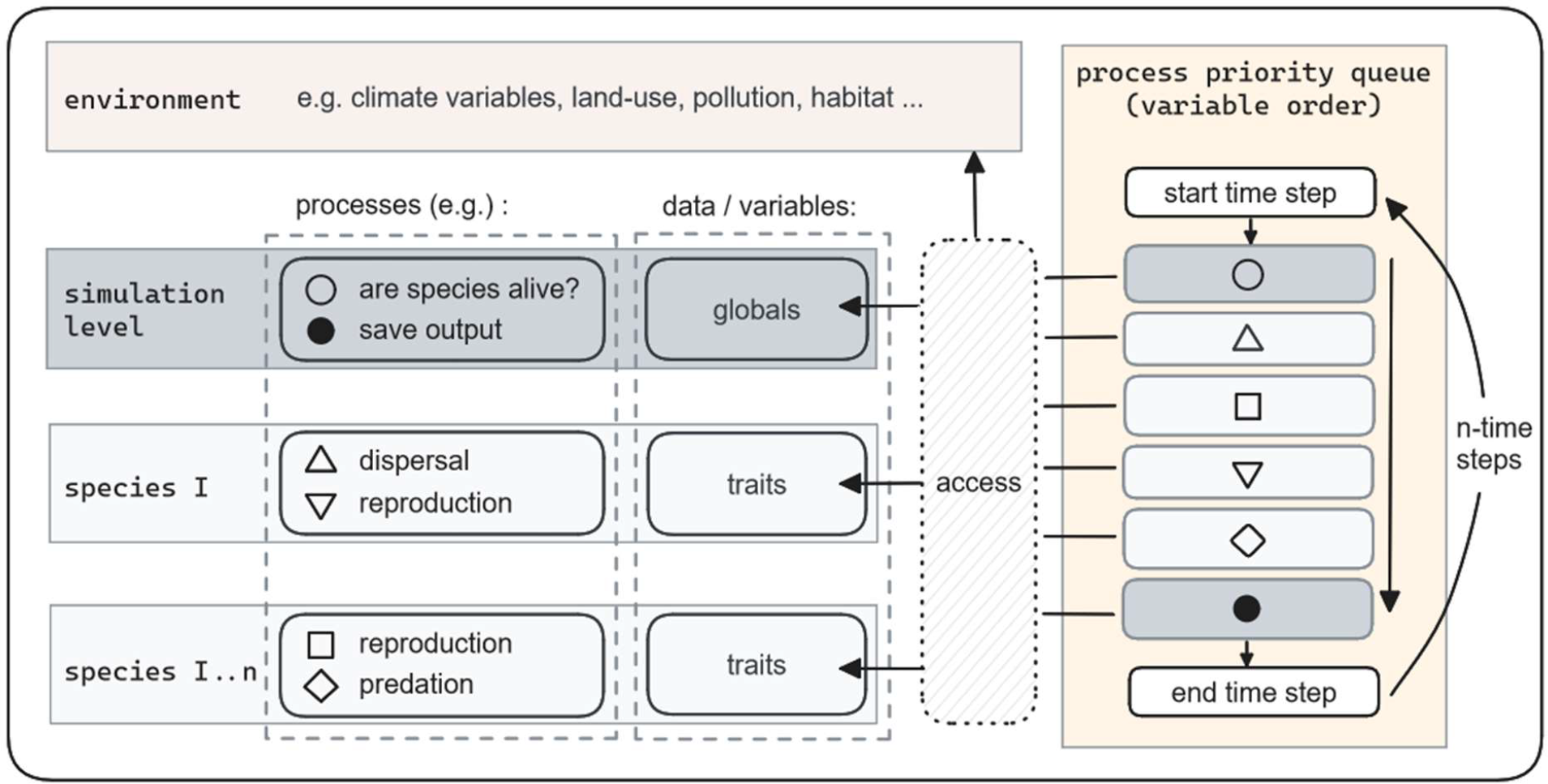
A high level overview of the objects in a metaRange simulation as well as their relation to each other. Note that a process may either be attached to a specific species or to the simulation itself (indicated by the symbols). These processes are executed each time step in a variable order based on the priority the user assigns to them (box: process priority queue). Each of the processes has access to the simulation environment, global variables (named “globals” in the package) and the traits of the species.

The simulation contains 1) an ‘environment’ that holds the relevant environmental factors (in form of raster data) and 2) one or more ‘species’ that are simulated in the environment. Each species has two main characteristics: ‘traits’, which are data describing attributes of the species, and ‘processes’, which are functions describing the species interaction with its surroundings (time, environment, other species, etc.).

Note that the traits are not only ‘functional traits’ (Violle et al., 2007), but use a wider definition that includes species-level state variables (e.g. is the species extinct or alive), hyperparameters (e.g. maximum dispersal distance), and population-level state variables (e.g. abundance).

The biological (e.g. reproduction, dispersal) and computational (e.g. saving output to a file) processes are executed in a user-defined order at each simulated time step. The processes can access and modify the species traits (e.g. abundance) and variables that are stored on a global simulation level (e.g. number of species in each cell).

To enable a quick start into the package, it includes functions that describe common ecological processes such as: climate suitability calculation, reproduction, dispersal, and metabolic scaling. Fig. 2 provides an example of what a simulation could look like. Because of the highly modular structure of the package the user can build a large variety of models with the functions already provided, or extend the package with user-defined functions. There is no limit on the number of environmental factors, species traits, or processes and only little computational requirement on the type of traits and processes that can be used. Note that a model built with metaRange does not simulate the environment, it only accesses the raster data supplied by the user as input (Fig. 2).

**Figure 2:**
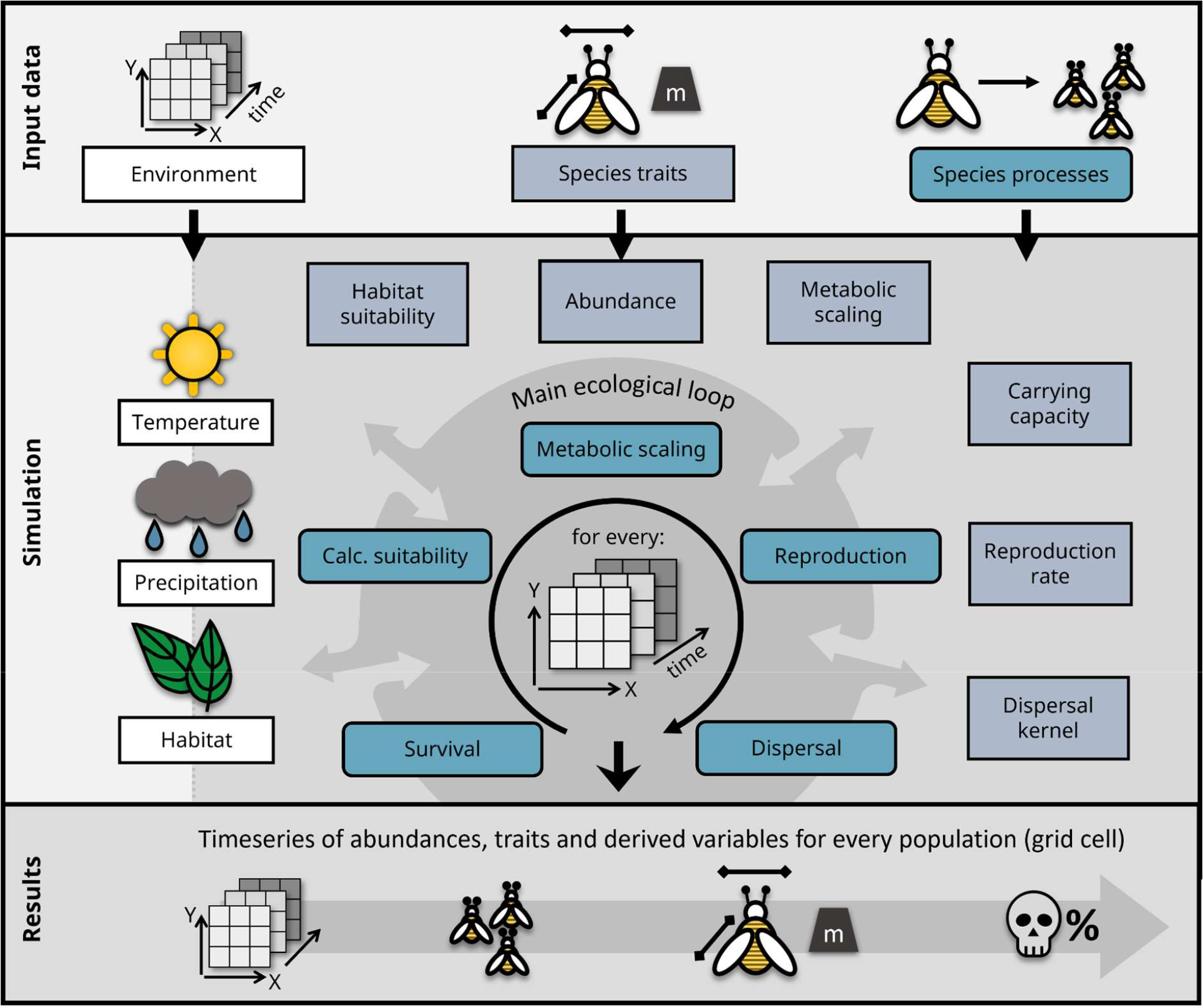
Example of some of the different environmental factors, species traits and processes that could be included in a simulation.

Below, we give a short overview over the functions that are already provided in metaRange v1.x.

### 3.1 Environmental suitability

The shape of the ecological response curve to a given variable (i.e. the species niche related to this variable) has been subject of debate (Hirzel and Le Lay, 2008; Oksanen and Minchin, 2002). The package provides a function that estimates this response curve as local environmental suitability based on a beta distribution, using the three “cardinal” values, i.e. the minimum (*V*_*min*_), optimum (*V*_*opt*_) and maximum (*V*_*max*_) of the species for a given environmental niche (*V*_*env*_). This function is based on a formula provided by Yin et al. (1995) and simplified by Yan and Hunt, (1999):

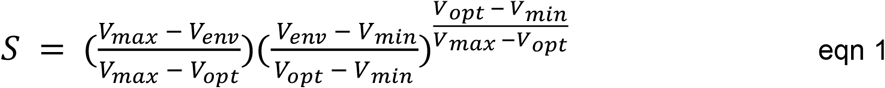

The beta distribution has the advantage that it can assume a variety of shapes, ranging from symmetric to skewed (Austin et al., 1994). Additionally, in this specific form of the function, all free parameters are biologically meaningful and could be measured through experiments. Note that there are many other response shapes of the suitability (Oksanen and Minchin, 2002), which the user could implement within the metaRange framework.

### 3.2 Metabolic scaling

A key metaRange feature is the possibility to use metabolic scaling based on the metabolic theory of ecology (MTE) (Brown et al., 2004) to constrain different species-specific parameters in relation to body temperature (local environmental temperature for plants and ectotherm animals) and body mass. The equation is split into three factors: a parameter- and species-specific normalization constant *Y*_0_, the mass *M* scaled with a parameter-specific allometric scaling exponent *a* and the Boltzmann factor that includes a parameter-specific activation energy *E*, the boltzmann constant *K*_*B*_ and the absolute body temperature *T*. The typical values for the different processes can be found in table 1.

**Table 1:**
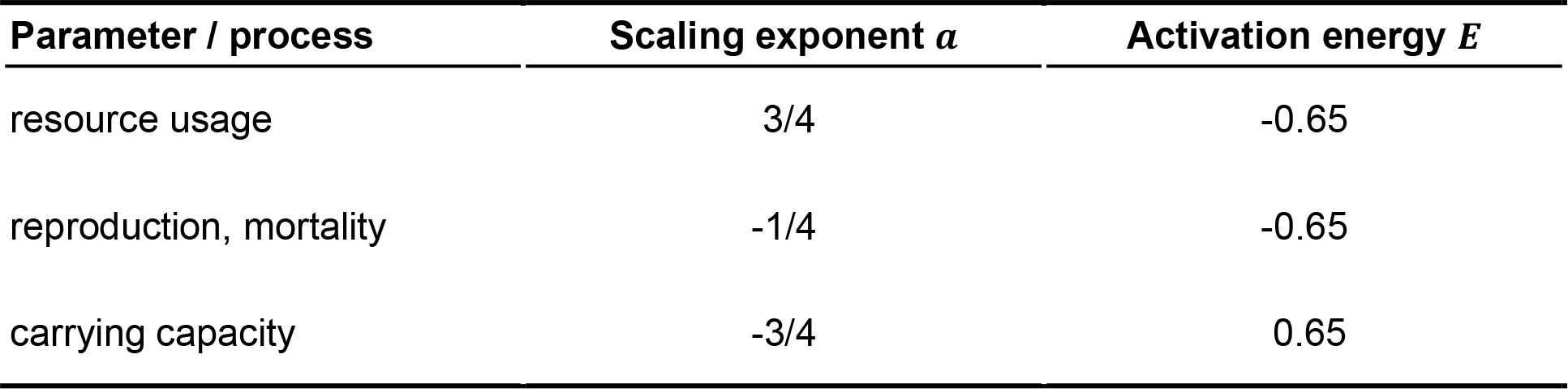
Commonly used values for the scaling exponent and the activation energy for different processes (adapted from table 2 in Brown et al., (2012)).

The MTE predicts that closely related species have similar *Y*_0_, which greatly reduces parametrization time while keeping a high-level of complexity.

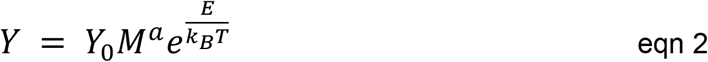

As most species lack experimentally measured values for *Y*_0_, the package includes a function to estimate *Y*_0_ based on parameter values under a reference temperature (See details in Supporting Information, S1).

### 3.3 Population dynamics

To simulate local population dynamics, metaRange features an implementation of the Ricker equation (Ricker, 1954):

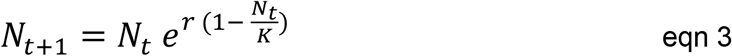

Where *N* is the number of individuals, *t* is a point in time, *r* is the per-capita reproduction rate, and *K* is the carrying capacity.

The package also includes a second version of the Ricker model that includes Allee effects (Cabral and Schurr, 2010):

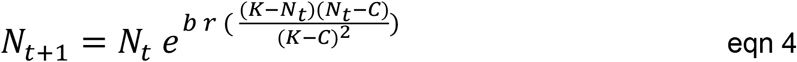

Where *C* is the Allee threshold and *b* is an unitless factor controlling the overcompensatory strength of the model.

### 3.4 Dispersal

Individuals are dispersed from a source cell via a dispersal kernel. If the abundance is described by a matrix ***N*** with the number of rows and columns *m, n*, then for a singular cell with the coordinates *i* ∈ ℤ : 1 ≤ *i* ≤ *m* and *j* ∈ ℤ : 1 ≤ *j* ≤ *n* this process is described by the following equation:

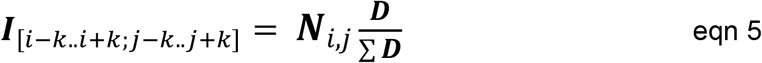

Where ***I*** is an intermediate matrix of the same size as ***N*** that stores the dispersing individuals and ***D*** is the dispersal kernel (a square matrix of uneven order). The term [*i* − *K*.. *i* + *K*; *j* − *K*.. *j* + *K*] describes the submatrix of ***I*** that is covered when the dispersal kernel is centered on the cell *i, j*. Consequently, *K* describes the radius of the dispersal kernel in grid cells (i.e. the maximum dispersal distance).

The algorithm uses reflexive boundary conditions (i.e. no individuals get lost at the edges of the landscape). The final number of dispersed individuals for each sink cell is the sum of all matrices ***I*** that are calculated for the source cells.

For many species, dispersal is more complex than a symmetrical kernel, involving environmental perception and even memory to generate directed movement towards a specific target, or influenced by other external factors. To allow this type of movement choice, the dispersal process can be influenced by “weights”, which are used to redistribute the individuals within the dispersal kernel, resulting in heterogeneous dispersal (Savary et al., 2023). If, for example, individuals are able to perceive habitat quality, a habitat suitability map can be used as weights, which results in individuals moving towards a more suitable habitat. This process can be expressed by expanding equation 5 to:

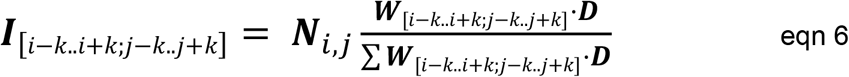

Where ***W*** is a matrix of weights, with the same size as ***N***.

By default, the kernel used a negative exponential distribution (Nathan et al., 2012):

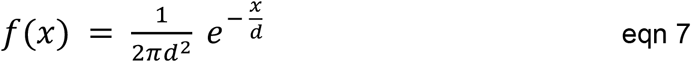

Where *x* is the distance to the source cell and *d* is the mean dispersal distance.

## 4 Example application

To highlight key features and illustrate usage, we conducted an experiment testing the common ecological concept of a generalist-specialist tradeoff.

### 4.1 Experimental design

We simulated 100 virtual grassland insect species under realistic environmental fluctuations in Germany on a 1 km^2^ resolution (> 500.000 grid cells) over 110 years (10 years to let the initial populations reach a quasi-equilibrium from initialization + 100 years simulation).

The species had the following processes scheduling: Calculation of the climate niche suitability (equation 1), metabolic scaling of the reproduction rate and carrying capacity (equation 2), adjustment of these values based on the habitat and climate suitability, reproduction described by the Ricker equation (equation 3), kernel-based weighted dispersal (equation 6 and 7), and, depending on the scenario, different forms of competition. A full model description can be found in Supporting Information S1.

The three competition scenarios we included were: A) No competition between the species, B) Competition between the species but without a generalist-specialist trade-off, C) Competition between the species with a strong generalist-specialist trade-off in which the maximum niche suitability of each species is inversely related to its niche width.

The two competition scenarios use a global carrying capacity as interaction currency (Kissling et al., 2012) given by the grid cell’s grassland cover.

### 4.2 Results

The runtime of each scenario was ∼40 minutes on a regular desktop computer (AMD Ryzen 7 5700G, 16GB RAM).

We find a difference in the regionwide number of surviving species (gamma richness) and in the maximum number of species in a grid cell (alpha richness) between the competition scenarios. In scenario A (no competition), 78 species survive, and the local richness is the highest (Fig. 3). Scenario B and C reveal a similar local richness, which is lower compared to scenario A, but differ in the gamma richness. In scenario B, 66 species survive, while in scenario C 75 species survive.

**Figure 3:**
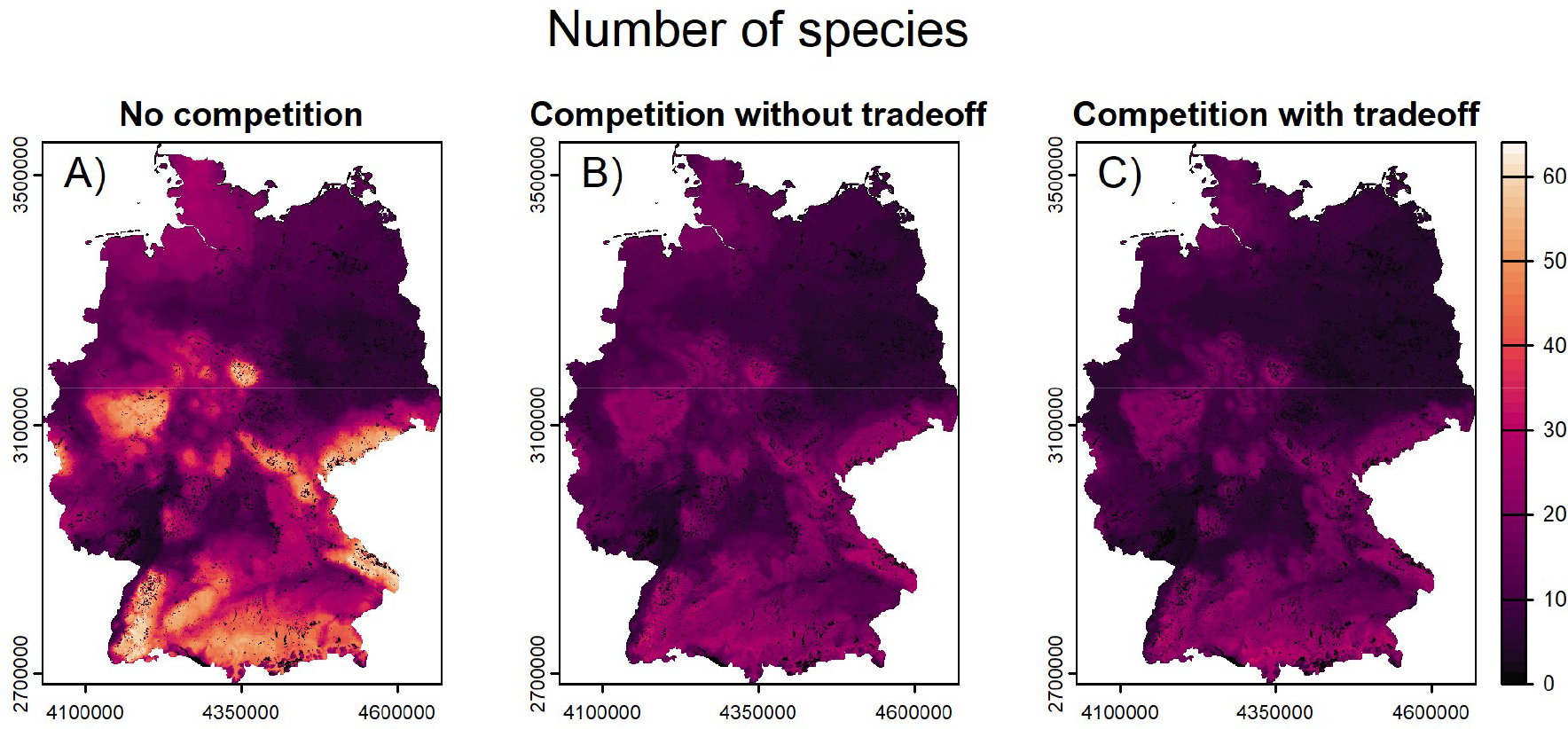
The aggregated number of living species (abundance > 1) in each cell at the last time step of the simulation. Note that the small black dots are cells without any grassland and therefore without any of the species.

To compare the effect of the generalist-specialist trade-off, we plot the percentage of occupied vs potentially suitable habitat (i.e. the range filling) of the species that survive in at least one of the three scenarios (Fig. 4). Compared to scenario A, the range filling of the species in the scenarios with competition is on average lower, but the niche width modulates the effect. In scenario B, the species with a narrow niche (i.e. specialists) lose more of their potential suitable range and many go extinct. In scenario C, the specialized species have a higher range filling compared to the generalist species, with fewer species going extinct.

**Figure 4:**
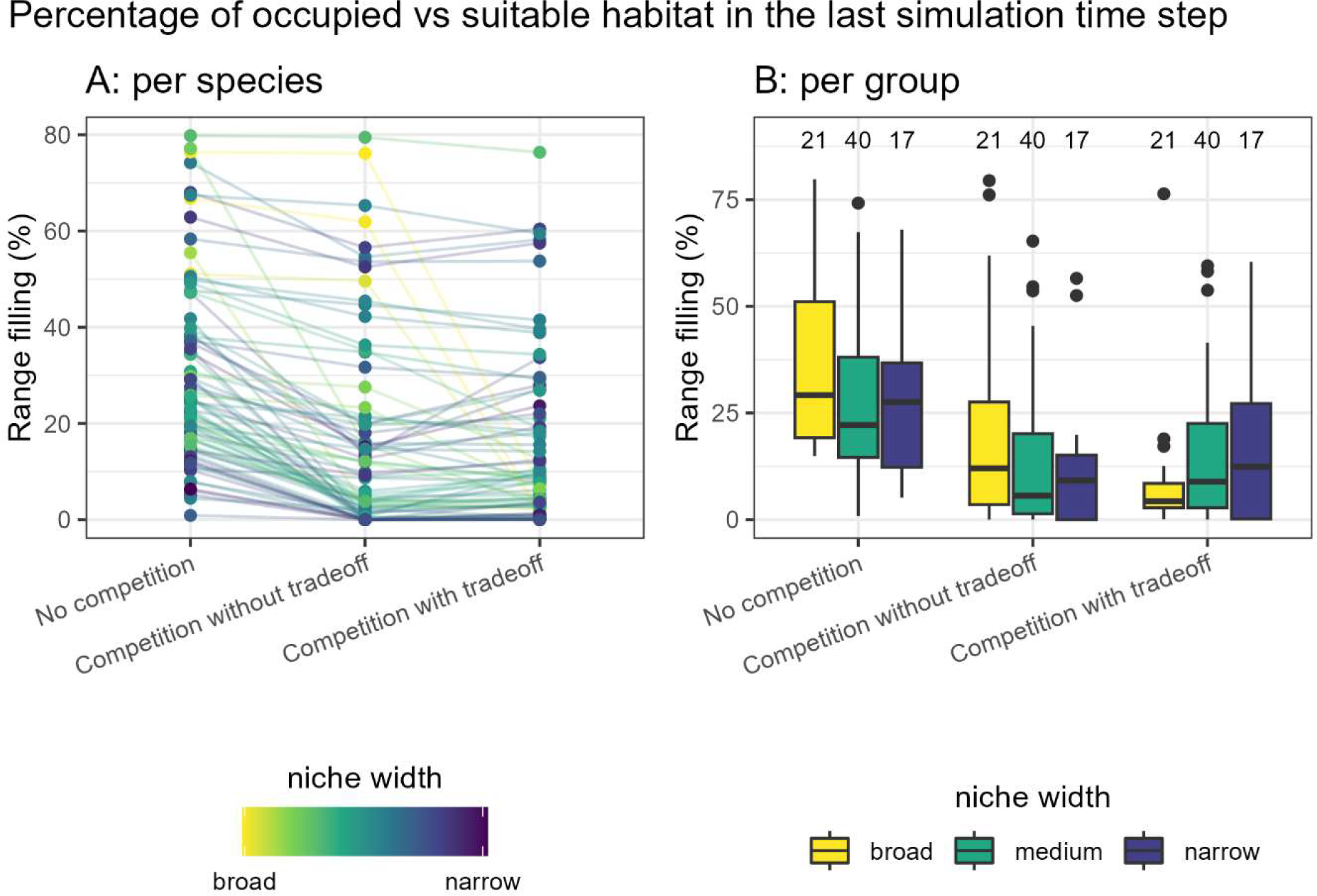
Range filling (occupied/suitable range) in the last simulation time step. A) Results per species. B) Results per groups with a broad (> 0.6), medium (0.4-0.6) or narrow (< 0.4) niche width. A niche width of 1 means that the species’ environmental limits equal the landscape’s environmental limits. Note that we do not perform statistical tests, as significance in simulation experiments can be reached simply by increasing replication.

## 5 Discussion

The result highlights how a fitness trade-off between specialist and generalist species aids the coexistence of these species under competition. Without this trade-off, the generalist species outcompete the specialist species, driving the latter to extinction. Hence, metaRange successfully emulates established ecological patterns (Wilson and Yoshimura, 1994).

Few previous mechanistic models also describe the range dynamics of species as for example: “rangeshiftR” (Bocedi et al., 2021, 2014; Malchow et al., 2020), “gen3sis” (Hagen et al., 2021) and “CATS” (Gattringer et al., 2023).

While they contain partially similar features, there remains a modelling gap combining the ability to simulate species interactions (e.g. rangeshifteR focuses on single species), ecological time scales (e.g. gen3sis focuses on evolutionary time scales) and generality to be applicable to various taxa (e.g. CATS focuses on plants).

Hence, metaRange fills this gap of high spatial resolution at range-wide extents and at ecological time scales of an interacting metacommunity of virtually any taxon and number of species.

## 6 Limitations & perspectives

Since all species share one rasterized environment, in which one grid cell represents the habitat of one population, species simulated together need to have a similar home range size for this assumption to be realistic. It should also be noted that in version 1.x of the package, the kernel-based dispersal does not include gene flow.

Nevertheless, the current version can already be adapted to a wide variety of study systems, while any missing processes can easily be introduced due to metaRange’s modularity.

## 7 Conclusions

The metaRange package offers the possibility to simulate a wide variety of species and species interactions with various environmental factors on a high-resolution landscape. Based on the number of species and processes included, only a comparatively small number of parameters need to be calibrated, enabling faster model development and hypotheses testing. We believe that metaRange’s modular structure and its focus on populations rather than individuals allows for wider applicability across organism groups and study questions.

## Acknowledgements

S.F. acknowledges funding by the “Deutsche Bundesstiftung Umwelt” (DBU). S.F. and J.S.C. acknowledge funding by the Bavarian Ministry of Science and Arts through the BLIZ project within the “Bayklif” network. Although this study is neither officially nor implicitly endorsed by the EU, we acknowledge that the Copernicus Land Monitoring Service products that are used in the example application were produced “with funding by the European Union”.

