## Supporting Information, S1 for "metaRange: A framework to build mechanistic range models"

### Overview, Design concepts, Details

#### Introduction

This is the model description of the example application from the manuscript "metaRange: A framework to build mechanistic range models". The description is based on the ODD (Overview, Design concepts, Details) protocol (Grimm et al., 2010, 2006). The data is archived under the DOI: (10.5281/zenodo.10792585).

#### Purpose & design

The purpose of this model is to illustrate how the metaRange R package can be used, exemplified with an experiment that tests the effects of the common ecological theory of a generalist-specialist tradeoff in the context of different levels of competition between the species.

##### Entities, state variables, and scales

Table 1 lists the environmental variables and table 2 lists the species variables used in the model.

*Table 1: Environment variables*

| Name | Scale & type / unit | Description |
|---|---|---|
| BIO10 | Germany, 1km2, 110 years; Kelvin | Bioclimatic variable 10 |
| BIO18 | Germany, 1km2, 110 years; millimeters | Bioclimatic variable 18 |
| grassland | Germany, 1km2, 110 years; 0-1 | Percentage of grassland / 100 |

*Table 2: Species variables*

| Trait name | Scale & type / unit | Description |
|---|---|---|
| abundance | 2D matrix; landscape extent; numeric | The number of individuals in each cell of the landscape |
| BIO10_maximum | scalar; numeric, Kelvin | Upper limit for bioclimatic variable 10 |
| BIO10_minimum | scalar; numeric, Kelvin | Lower limit for bioclimatic variable 10 |
| BIO10_optimum | scalar; numeric, Kelvin | Preferred value of bioclimatic variable 10 |
| BIO18_maximum | scalar; numeric, millimeters | Upper limit for bioclimatic variable 18 |
| BIO18_minimum | scalar; numeric, millimeters | Lower limit for bioclimatic variable 18 |
| BIO18_optimum | scalar; numeric, millimeters | Preferred value of bioclimatic variable 18 |
| carrying_capacity | 2D matrix; landscape extent; numeric | Maximum number of individuals per population |
| carrying_capacity_mte_constant | 2D matrix; landscape extent; numeric | The constant used in the MTE to scale the carrying capacity |

| | | |
|---|---|---|
| dispersal_kernel | 2D matrix; extent = 2 * maximum_dispersal_dist + 1; numeric | The dispersal kernel used to disperse the populations during the dispersal process |
| exponent_carrying_capacity | scalar; numeric | The exponent used in the MTE to scale the carrying capacity |
| exponent_reproduction_rate | scalar; numeric | The exponent used in the MTE to scale the carrying capacity |
| mass | 2D matrix; landscape extent; numeric; gram | The mean body mass of an individual in each population |
| maximum_dispersal_dist | scalar; integer; km | The maximum dispersal distance |
| mean_dispersal_distance | scalar; numeric; km | The mean dispersal distance |
| niche_width | scalar; numeric; 0-1 | The percentage of the mean of the range between both the climatic variables in relation to the range of the environmental values of the whole landscape |
| reproduction_rate | 2D matrix; landscape extent; numeric | The per-capita reproduction rate |
| reproduction_rate_mte_constant | scalar; numeric | The constant used in the MTE to scale the reproduction rate |
| suitability | 2D matrix; landscape extent; numeric; 0-1 | The climate suitability of each grid cell for the population living there |

### Process overview and scheduling

Following is a pseudocode description of the model logic:

```
for each time step:
│   // Process 1
│   for species in alive_species:
│   │   species[suitability] =
│   │   │   calculate_suitability(
│   │   │   │   vmax = species[BIO10_maximum],
│   │   │   │   vopt = species[BIO10_optimum],
│   │   │   │   vmin = species[BIO10_minimum],
│   │   │   │   venv = species[environment[BIO10]
│   │   │   ) *
│   │   │   calculate_suitability(
│   │   │   │   vmax = species[BIO18_maximum],
│   │   │   │   vopt = species[BIO18_optimum],
│   │   │   │   vmin = species[BIO18_minimum],
│   │   │   │   venv = environment[BIO18])
│   │   │   species[suitability] < suitability_cutoff = 0
│   │   if scenario == "competition_with_tradeoff":
│   │   │   species[suitability] = species[suitability] * (1 - species[niche_width])
│   // Process 2
│   for species in alive_species:
│   │   species[reproduction_rate] =
│   │   │   metabolic_scaling(
│   │   │   │   normalization_constant = species[reproduction_rate_mte_constant],
│   │   │   │   scaling_exponent = species[exponent_reproduction_rate],
│   │   │   │   mass = species[mass],
│   │   │   │   temperature = environment[BIO10],
│   │   │   │   E = -0.65,
│   │   │   │   k = 8.617333e-05
│   │   │   ) * species[suitability]
│   │   species[carrying_capacity] =
│   │   │   metabolic_scaling(
│   │   │   │   normalization_constant = species[carrying_capacity_mte_constant],
│   │   │   │   scaling_exponent = species[exponent_carrying_capacity],
│   │   │   │   mass = species[mass],
│   │   │   │   temperature = environment[BIO10],
│   │   │   │   E = 0.65,
│   │   │   │   k = 8.617333e-05
│   │   │   ) * environment[grassland] * species[suitability]
│   │   truncate species[carrying_capacity]
│   // Process 3
│   for species in alive_species:
│   │   species[abundance] =
│   │   │   ricker_reproduction_model(
│   │   │   │   species[abundance],
│   │   │   │   species[reproduction_rate],
│   │   │   │   species[carrying_capacity]
│   │   │   )
│   // Process 4
│   for species in alive_species:
│   │   species[abundance] =
│   │   │   dispersal(
│   │   │   │   abundance = species[abundance],
│   │   │   │   weights = species[suitability] * environment[grassland],
│   │   │   │   dispersal_kernel = species[dispersal_kernel]
```

```
|   |   |   )
|   // Process 5
|   if scenario == "competition_with_tradeoff" or "competition_without_tradeoff":
|   |   for species in alive_species:
|   |   |   sum_abundance = sum_abundance + species[abundance]
|   |   competition_pressure = sum_abundance / global_carrying_capacity
|   |   clamp competition_pressure to [0, 1]
|   |   for species in alive_species:
|   |   |   species[abundance] =
|   |   |   |   species[abundance] * (1 - competition_pressure) +
|   |   |   |   species[abundance] * species[suitability] * competition_pressure
|   |   |   truncate species[abundance]
|   // Process 6
|   for species in alive_species:
|   |   current_abundance = sum of abundance
|   |   append current_abundance to abundance_list[species]
|   |   if current_abundance < 1:
|   |   |   remove species from alive_species
|   |   suitable_cells = sum of suitability > 0
|   |   append suitable_cells to suitable_cells_list[species]
|   |   occupied_cells = number of cells where abundance > 1
|   |   append occupied_cells to occupied_cells_list[species]
// Process 7
save results
```

### Design concepts

#### Basic principles

The simulation contains an `environment` that holds and manages all the environmental factors that may influence the simulation (in form of raster data) and it contains multiple `species` that are simulated in the environment. Each species has two main characteristics: `traits` which are pieces of data that describe attributes of the species and `processes` which are functions that describe the species interaction with its surroundings (time, climate, other species, etc.).

Note that the traits are not only `functional traits` as defined by (Violle et al., 2007), but use a wider definition that can include species-level state variables (e.g. is the species extinct or alive), hyperparameter (e.g. maximum dispersal distance) and population-level state variables (e.g. abundance).

Species have implied populations, each of which inhabit one grid cell of the landscape. On a computational level this means that the traits of a species will in most cases be stored in a matrix with the same size as the landscape it is simulated in, where each value in the matrix represents the trait value of a population.

**Emergent properties**

Emergent properties of the model are the local abundances and traits of the populations, the metapopulation dynamics, the overall distribution of the species in the landscape, the range filling (potential range vs realized range) as well as the long-term survival.

**Adaptation & Sensing**

Through the use of weighted dispersal, the implicit individuals are moving towards a more suitable habitat during dispersal, if they have the ability to perceive and reach it. The reproduction rate and the carrying capacity are scaled according to the MTE and adjusted based on the climate suitability and habitat suitability.

**Interaction**

The model includes three scenarios: A) No competition between the species, B) Competition between the species but without a generalist-specialist trade-off, C) Competition between the species with a strong generalist-specialist trade-off in which the maximum niche suitability of each species is inversely related to its niche width (I.e. the range that their environmental limits cover compared to the range of the environmental values in the landscape).

The two competition scenarios use indirect competition for resources based on a "global carrying capacity" as interaction currency (Kissling et al., 2012), which is calculated from the amount of grassland in each habitat cell.

**Observation**

The data used in the analysis are the aggregated number of living species (abundance > 1) in each cell at the last time step of the simulation and the percentage of occupied vs potentially suitable habitat (range filling) in the last simulation time step as well as trait data from the species (niche width). This data is considered in its entirety, without any sampling.

#### Initialization & input data

The initialization was the same for all three scenarios. The mass and environmental limits of species were randomly selected (but using the same seed, therefore the same species were used in each scenario), while the other traits were set to fixed estimated values. An overview of the trait values for all species can be found in table 3.

*Table 3: Initial values*

| Trait name | Initial value | Notes |
|---|---|---|
| abundance | 2000 | - |
| BIO10_maximum | random | uniform drawn between the mean and max of the environment BIO10 |
| BIO10_minimum | random | uniform drawn between the mean and min of the environment BIO10 |
| BIO10_optimum | random | uniform drawn between the BIO10_maximum and BIO10_minimum |
| BIO18_maximum | random | uniform drawn between the mean and max of the environment BIO18 |
| BIO18_minimum | random | uniform drawn between the mean and min of the environment BIO18 |
| BIO18_optimum | random | uniform drawn between the BIO18_maximum and BIO18_minimum |
| carrying_capacity | 100000 | - |
| carrying_capacity_mte_constant | derived | calculated using the function `calculate_normalization_constant` with: parameter_value = carrying_capacity, scaling_exponent = exponent_carrying_capacity, mass = mass, reference_temperature = BIO10_optimum, E = 0.65, k = $k_B$ |
| dispersal_kernel | derived | Calculated from maximum_dispersal_dist and mean_dispersal_distance with a negative exponential function |
| exponent_carrying_capacity | -3 / 4 | (Brown et al., 2012) |
| exponent_reproduction_rate | -1 / 4 | (Brown et al., 2012) |
| mass | random | uniform drawn between [0.5, 5] |

| | | |
|---|---|---|
| maximum_dispersal_dist | derived | ceiling of mass * 2 |
| mean_dispersal_distance | derived | mass / 3 |
| niche_width | derived | The percentage of the mean of the range between both the climatic variables in relation to the range of the environmental values of the whole landscape |
| reproduction_rate | 0.5 | - |
| reproduction_rate_mte_constant | derived | calculated using the function `calculate_normalization_constant` with: parameter_value = reproduction_rate, scaling_exponent = exponent_reproduction_rate, mass = mass, reference_temperature = BIO10_optimum, E = -0.65, k = $k_B$ |
| suitability | - | Calculated when model runs |

#### Input data

The metaRange package doesn't generate the environment and time series in which the simulation takes place, it only accesses previously generated raster data. In this example model, we used the percentage of grassland (Copernicus Land Monitoring Service et al., 2018) to restrict the maximal carrying capacity of the habitat and the bioclimatic variables (Karger et al., 2018, 2017) 10 (mean daily mean air temperatures of the warmest quarter) and 18 (mean monthly precipitation amount of the warmest quarter) as environmental niche variables. To generate a realistic time-series with auto-correlated fluctuations, we used reference climate data from the German Meteorological Service from the period 1901-2000 (Deutscher Wetterdienst, 2024). We centered these timeseries and applied them as an offset to each year in the simulation (excluding the burn-in phase).

### Submodels

#### Environmental suitability

The shape of the ecological response curve to a given variable (i.e. the species niche related to this variable) has been subject of debate (Hirzel and Le Lay, 2008; Oksanen and Minchin, 2002). We used a function provided by metaRange "calculate_suitability" that can estimate the local environmental suitability for a given environmental value, based on a beta distribution, using the three "cardinal" values, i.e. the minimum ($V_{min}$), optimum ($V_{opt}$) and maximum ($V_{max}$) of the species for that environmental niche ($V_{env}$). This function is based on a formula provided by Yin et al., (1995) and simplified by Yan and Hunt, (1999):

$$S_{i,j} = \left(\frac{V_{max} - V_{env_{i,j}}}{V_{max} - V_{opt}}\right)\left(\frac{V_{env_{i,j}} - V_{min}}{V_{opt} - V_{min}}\right)^{\frac{V_{opt} - V_{min}}{V_{max} - V_{opt}}} \quad \text{eqn 1}$$

The beta function has generally the benefit that it can assume a variety of shapes, ranging from symmetric to skewed (Austin et al., 1994). Additionally, in this specific form of the function, all free parameters are biologically meaningful and could be measured through experiments, thus increasing the mechanistic link between physiological traits and emergent patterns.

To calculate the climate suitability, we separately calculate the suitability for the environmental variables (BIO10, BIO18) and multiplied them.

**Metabolic scaling**

We used the metaRange function "metabolic_scaling" that uses the metabolic theory of ecology (MTE) (Brown et al., 2004) to scale the reproduction rate and the carrying capacity of each species, in relation to the temperature in the landscape and the average mass of the individuals of the species. The equation is split into three factors: a parameter- & species-specific normalization constant $Y_0$, the mass $M$ scaled with a parameter specific allometric scaling exponent $a$ and the Boltzmann factor that includes a parameter specific activation energy $E$, the boltzmann constant $k_B$ and the absolute temperature $T$.

$$Y_{i,j} = Y_0 M^a e^{\frac{E}{k_B T_{i,j}}} \quad \text{eqn 2}$$

The typical values for the different processes can be found in table 4.

*Table 4: Commonly used literature values for the scaling exponent and the activation energy for different processes. This is adapted from table 2 in Brown et al. (2012).*

| Parameter / process | Scaling exponent $a$ | Activation energy $E$ |
|---|---|---|
| resource usage | 3/4 | -0.65 |
| reproduction, mortality | -1/4 | -0.65 |
| carrying capacity | -3/4 | 0.65 |

We used the function “calculate_normalization_constant” to estimate the normalization constant needed for the metabolic scaling, based on the input values for the reproduction rate and the carrying capacity (table 3). This function is inverse to equation 2 defined as:

$$Y_0 = \frac{Y}{M^a e^{\frac{E}{k_B T}}} \qquad \text{eqn 3}$$

The reference temperature $T$ was set to the temperature optimum of the species (BIO10_optimum). After the metabolic scaling we multiplied the reproduction rate with the climate suitability, and we multiplied the carrying capacity with both the climate and the habitat suitability (i.e. the percentage of available grassland).

**Population dynamics**

To simulation local population dynamics, we use the metaRange function “ricker_reproduction_model” that features an implementation of the Ricker reproduction model (Ricker, 1954) in the form of:

$$N_{t+1,i,j} = N_{t,i,j}\, e^{r_{t,i,j}\,(1-\frac{N_{t,i,j}}{K_{t,i,j}})} \qquad \text{eqn 4}$$

Where $N$ is the number of individuals in a population, $t$ is a point in time, $r$ is the per-capita reproduction rate and $K$ is the carrying capacity of the habitat.

**Dispersal**

We used the metaRange function "dispersal" in which the dispersal is modeled as a dispersal kernel that is applied to each cell of the abundance matrix and that distributes the individuals from a source cell to the surrounding cells, influenced by weights (habitat suitability multiplied with the climate suitability), which are used to redistribute the individuals within the dispersal kernel, resulting in heterogeneous dispersal (Savary et al., 2023). If the abundance is described by a matrix $\boldsymbol{N}$ with the number of rows and columns $m, n$, then for a singular cell with the coordinates $i \in \mathbb{Z} : 1 \leq i \leq m$ and $j \in \mathbb{Z} : 1 \leq j \leq n$ this process is described by the following equation:

$$\boldsymbol{I}_{[i-k..i+k;j-k..j+k]} = \boldsymbol{N}_{i,j} \frac{\boldsymbol{W}_{[i-k..i+k;j-k..j+k]} \cdot \boldsymbol{D}}{\sum \boldsymbol{W}_{[i-k..i+k;j-k..j+k]} \cdot \boldsymbol{D}} \quad \text{eqn 5}$$

Here, $\boldsymbol{I}$ is an intermediate matrix of the same size as $\boldsymbol{N}$, that stores the dispersing individuals from the source cell, $\boldsymbol{W}$ is a matrix of weights that has the same size as the abundance matrix which are used to redistribute the individuals within the dispersal kernel and $\boldsymbol{D}$ is the dispersal kernel (a square matrix of uneven order). The term $[i-k..i+k; j-k..j+k]$ describes the submatrix of $\boldsymbol{I}$ and $\boldsymbol{W}$ that is covered when the dispersal kernel is centered on the cell $i, j$. Consequently $k$ describes the radius of the dispersal kernel in grid cells (i.e. the maximum dispersal distance).

Note that this algorithm uses reflexive boundary conditions, which means that no individuals get lost at the edges of the landscape. This leads to a slightly higher density of individuals at the edges, though this effect is negligible on a sufficiently large landscape compared to the size of the kernel. This form of dispersal emulates environmental perception of the species to generate directed movement towards a specific target, in this case towards a more suitable habitat.

The result of the dispersal for each cell in the abundance matrix is then the sum of all the individual matrices $\boldsymbol{I}$ that are calculated for each grid cell.

We used the metaRange function "negative_exponential_function" to calculate the kernel values. This function uses the equation:

$$f(x) = \frac{1}{2\pi d^2} e^{-\frac{x}{d}} \qquad \text{eqn 6}$$

Where $x$ is the distance to the source cell and $d$ is the mean dispersal distance.

**Competition**

The competition is calculated through multiple steps. First, we calculate the overall number of individuals in each cell, i.e. sum of the abundances of the individual species $n$.

$$\text{sum}(\boldsymbol{N}) = \boldsymbol{N}_{1,i,j} + \boldsymbol{N}_{2,i,j} + \cdots + \boldsymbol{N}_{n,i,j} \qquad \text{eqn 7}$$

In the next step we calculate the competition pressure $\boldsymbol{P}$, which is the ratio of the sum of the abundances to the global carrying capacity $\boldsymbol{G}$, clamped to the range of zero to one.

$$P = \min(\max(\frac{\text{sum}(\boldsymbol{N})}{\boldsymbol{G}}, 0), 1) \qquad \text{eqn 8}$$

Lastly, we calculate the new abundance $N'$ for each species by computing a weighted mean between the abundance and the abundance multiplied with the suitability, weighted by the competition pressure.

$$N'_{n,i,j} = \lfloor N_{n,i,j}(1 - P_{i,j}) + N_{n,i,j}S_{n,i,j}P_{i,j} \rfloor \quad \text{eqn 9}$$

### Appendix

#### Technical details

The model was implemented in R version 4.3.2 (R Core Team, 2023) and is dependent on the following R packages: “terra” (Hijmans, 2023) is used for the geographical data handling, “R6” (Chang, 2021) is the underlying package that metaRange uses to implement the object-oriented structure, “Rcpp” (Eddelbuettel, 2013; Eddelbuettel et al., 2023; Eddelbuettel and Balamuta, 2018; Eddelbuettel and François, 2011) and “RcppArmadillo” (Eddelbuettel and Sanderson, 2014) are the packages that allows for the performance critical functions of metaRange to be implemented in C++, and “checkmate” (Lang, 2017) is used to ensure code stability.

#### Symbols

In table 5 you may find an overview of the different symbols in this manuscript, including the process in which they are used.

*Table 5: List of symbols used in this paper.*

| Symbol | Description | Usage | Scale / unit |
|---|---|---|---|
| $V_{max}$ | upper (i.e. maximum) tolerable value | suitability calculation | - |
| $V_{min}$ | optimal (i.e. preferred) value | suitability calculation | - |
| $V_{opt}$ | lower (i.e. minimum) tolerable value | suitability calculation | - |
| $V_{env}$ | environmental value for which to calculate the suitability | suitability calculation | - |
| $Y$ | parameter | metabolic scaling | - |
| $Y_0$ | Species and process specific normalization constant | metabolic scaling | - |
| $M$ | mean (individual) mass | metabolic scaling | gram |
| $a$ | allometric scaling exponent of the mass | metabolic scaling | - |
| $E$ | activation energy | metabolic scaling | electronvolts |
| $k_B$ | boltzmann constant | metabolic scaling | 8.617333e-05 |
| $T$ | absolute temperature | metabolic scaling | Kelvin |
| $r$ | per-capita reproduction rate | Ricker reproduction model | - |
| $N$ | abundance | Ricker reproduction model, dispersal | >0 |
| $K$ | carrying capacity | Ricker reproduction model | >0 |
| $d$ | mean dispersal distance | dispersal | >0 |
| $x$ | distance | dispersal | >0 |
| $D$ | dispersal kernel | dispersal | - |
| $W$ | weights | dispersal | - |

| | | | |
|---|---|---|---|
| $P$ | competition pressure | competition | 0-1 |
| $G$ | global carrying capacity | competition | 0-1 |
| $S$ | suitability | suitability calculation | 0-1 |
| $k$ | radius of the dispersal kernel in grid cells | dispersal | |
| $m, n$ | number of rows and columns of the traits matrices (including abundance) as well as the landscape raster | universal | -<br>$m, n \in \mathbb{Z}$ |
| $i, j$ | rows and column indices | universal | $i, j \in \mathbb{Z}$<br>$i : 1 \leq i \leq m$<br>$j : 1 \leq j \leq n$ |
